# Benefits of music training for learning to read: Evidence from cortical tracking of speech in children

**DOI:** 10.1101/2025.09.05.674218

**Authors:** Maria C. Garcia-de-Soria, Brian Mathias, Anne Keitel, Anastasia Klimovich-Gray

## Abstract

Musical training has long been argued to boost early phonological and reading abilities. Cortical tracking of speech (CTS) has been proposed as a mechanism for this music-to-literacy transfer. In this study, we examined how musical training shapes CTS in young readers and whether it facilitates literacy benefits. In a sample of 57 children aged 5-9, musical training was linked to enhanced reading and phonological awareness (PA). EEG during story listening revealed that higher left-hemispheric and lower right-hemispheric CTS were also associated with higher reading scores. However, children with higher musicality exhibited stronger reading skills at lower levels of left-hemispheric CTS, suggesting more adult-like speech analysis. Critically, PA mediated the relationship between musicality and reading: greater musicality was associated with stronger PA, which in turn predicted higher reading performance, independent of demographic and cognitive factors. These findings indicate that musical training supports literacy by enhancing PA and shaping left-lateralized speech processing.

## Introduction

The benefits of music training on cognitive development, particularly reading acquisition, have become a prominent area of research. Numerous studies have linked musical training with improved academic performance, including enhanced reading skills in school aged children^1,2,3^. The early years of formal education between the ages of 5 and 9 represent a critical period for significant cognitive and neural changes underlying the transition from emergent to proficient reading^4^. This is supported by the maturation of key neural pathways involved in speech processing, particularly the arcuate fasciculus, which connects the temporal and frontal language areas^5^, and the increased specialisation of the ventral stream, including the ventral occipitotemporal cortex, for letter processing during this time^6^. As children become more proficient readers, brain responses to complex language processing within the bilateral frontotemporal language network become increasingly more left-lateralized which is thought to reflect increasing specialisation to language-related tasks^7,8,9,10,11^. Therefore, early school years education is the optimal period for examining how musical training boosts literacy skills and neuro-cognitive mechanisms involved in language processing and reading acquisition.

### Understanding the Relationship Between Musical Training and Reading

While it is well-known that musical training positively influences reading development^12,13,14,15^ with long-lasting positive effects^3^, there remains a gap in understanding the specific neuro-cognitive pathways through which these benefits manifest, particularly during childhood. Several theoretical frameworks have been proposed to explain the transfer effects of musical training on language and literacy. The OPERA hypothesis^16^ suggests that the demands of music on the auditory system enhance low-level auditory encoding in overlapping speech-related circuits. Similarly, the PRISM model^17^ provides a mechanistic account of how rhythm-based musical activities foster shared auditory and sensorimotor neural circuits involved in both music and speech, including precision auditory analysis, temporal tracking of auditory stimuli, and the integration of auditory and motor responses.

These frameworks collectively emphasize that musical training has the potential to support language development through shared cognitive and neural pathways, particularly those involved in temporal synchronisation of neural systems to the external auditory information which benefits phonological processing and subsequently reading. This is consistent with behavioural evidence suggesting that musical training can strengthen auditory processing and phonological awareness (PA) – the ability to identify and manipulate the sound structure of language^17,18,19^—and enhance visual-motor coordination and timing skills^20,21^. Musical training has also been associated with improved performance in tasks involving fine-grained pitch discrimination and speech-in-noise perception^12,22,23^ and with greater longitudinal neural plasticity and flexibility within auditory networks^24^. Altogether, these findings suggest that musical training fosters a more adaptive auditory system, potentially making it better suited to process the complex and variable acoustic signals encountered in natural speech. By enhancing auditory precision and the coordination between auditory and higher-order cognitive systems, musical training may lay a stronger foundation for phonological awareness and, by extension, reading acquisition.

However, it is important to acknowledge that the evidence for far-transfer effects from music to language and reading remains highly contentious. Despite the compelling empirical findings and theoretical frameworks, a growing body of recent research has failed to replicate robust far-transfer effects, particularly in rigorously controlled trials. Schellenberg & Lima (2024)^25^ argue that many reported links between musical training and cognitive or academic outcomes are likely driven by pre-existing individual differences (e.g. socio-economic status, cognitive ability, parental involvement) rather than genuine causal transfer effects, which often fail to replicate under rigorous scrutiny. This debate has intensified in recent years, prompting for more cautious interpretations, especially for effects far from the trained domain^26^. While the current study builds on established neurocognitive frameworks proposing shared mechanisms between music training and reading, it also aims to critically evaluate the robustness of these associations under controlled conditions and explicitly assess the contribution of other cognitive and socio-economic factors. In doing so, we aim to clarify the potential (and limitations) of music-based interventions in supporting reading development.

### Cortical Tacking of Speech and Music Training

Recent research has turned to exploring specific neural mechanisms to explain how enhanced auditory processing may support language development. One such mechanism is Cortical Tracking of Speech (CTS)^27^, also referred to as neural entrainment or neural coupling—the brain’s ability to synchronise its neural activity to the rhythmic structure of speech^28,29^. This process involves the brain aligning its oscillatory neural activity to the amplitude of the envelope and rhythmic patterns of speech (primarily prosodic and syllabic), facilitating the segmentation of continuous speech into meaningful units. This process is thought to create optimal time-windows for fine-grain spectro-temporal analysis critical for phoneme processing.

While this theoretical account has been highly influential^27,30^, recent work has raised questions about whether this type of neural alignment directly supports language processing in the way originally proposed. For example, new evidence suggests that cortical tracking may reflect multiple overlapping processes (not all of which directly related to linguistic comprehension) and that the link between entrainment and phoneme-level processing is less direct than assumed. Atanasova et al. (2025)^31^ emphasize that despite the widespread influence of oscillation-based theories of speech processing, there is currently little unambiguous empirical evidence that endogenous brain rhythms are involved in speech tracking in the way these theories predict. Similarly, Lalor and Nidiffer (2025)^32^ argue that CTS is better explained by the summation of evoked responses rather than by oscillatory entrainment, and they suggest that the role of endogenous oscillations in natural speech processing has likely been overstated. Extending this critical perspective, Whiteford et al. (2025)^33^ failed to replicate previously reported links between musically trained adults and enhanced early auditory encoding (measured as frequency following response), further questioning strong claims about the causal role of rhythmic neural alignment and experience-dependent plasticity in speech processing.

Despite these challenges, CTS has been shown to be linked to phonological awareness and play a key role in reading acquisition^34,35,36,37^. Previous research has demonstrated that both children and adults with dyslexia exhibit atypical neural tracking of speech rhythms, particularly in the delta (1–3 Hz) and theta (4–8 Hz) frequency bands corresponding to the analysis of prosodic and syllabic structure of speech^38^. These deficits in cortical tracking are thought to lead to difficulties with phonological processing over time that are central to dyslexia and other developmental language disorders^37^. Musical training is proposed as a potential intervention tool to enhance CTS by strengthening the auditory system’s sensitivity to temporal patterns in speech^39^.

While many studies have examined cortical tracking of separate features of speech, few have investigated the tracking of continuous, natural speech in children. Recently, Rogachëv and Sysoeva (2024)^40^ found a strong correlation between cortical tracking of natural speech and receptive speech abilities. Specifically, children aged 3 to 8 who exhibited stronger neural synchronization to natural speech—as measured by electroencephalogram (EEG)—also showed higher scores on standardized measures of speech comprehension and vocabulary understanding. Notably, acoustic tracking was associated with activity in right temporal regions, whereas semantic tracking engaged left fronto-central and right parieto-occipital regions. These findings suggest that cortical tracking of natural speech may serve as a neural marker of receptive language development in early childhood.

Previous findings suggest that musical training may enhance CTS and in doing so reinforce the neural mechanisms that support phonological processing and reading development^12,13,39^. In line with this, musical training has been shown to improve PA and reading skills by strengthening the brain’s ability to process rhythmic and temporal patterns^16,34^. However, most of related studies have focused on cortical tracking of rhythmic structures or isolated syllables, with no current research known to authors exploring how musicality simultaneously affects cortical tracking of natural speech, phonological skills and reading development in young children.

### The Current Study

In the present study we used EEG to investigate the relationship between reading, phonological skills, cortical tracking of continuous speech and musical engagement, in children undergoing music education beyond their standard school curriculum. Specifically, we assessed CTS (using mutual information analyses), PA, and musical training as potential predictors of reading success, while controlling for demographic and behavioural factors known to influence literacy outcomes, including socio-economic status, domain-general cognitive skills, and rhythmic motor coordination. Musical training was assessed through a caregiver-report questionnaire, which captured whether the child had received formal musical instruction and detailed their level of engagement in musical activities (e.g., hours spent playing an instrument). To our knowledge, this is the first study to provide a comprehensive review of the links between musical training, CTS and reading development, using neural processing of continuous natural speech as a core methodological approach in young children.

We had four hypotheses: 1) Musically trained children were expected to outperform the control group in literacy-related abilities, including PA, reading accuracy, and rhythmic tapping, consistent with previous findings on the link between musical training and language skills^13,14^. 2) We predicted that children with musical training would show stronger CTS, reflecting advanced abilities of speech signal perceptual analysis. Furthermore, we expected that this neural tracking would be more left-lateralized (as patterns observed in older children and adults) as a signature of linguistic maturation^8,9,10^. 3) We hypothesized that musical training and CTS would emerge as significant predictors of reading ability, having controlled for other non-linguistic behavioural and demographic factors. 4) Finally, we hypothesized that PA would mediate the relationship between musical training and reading ability, consistent with previous proposals that link musical training to enhanced perceptual analyses and parsing of auditory inputs^13,41^. Additionally, we proposed that CTS measures (particularly within the left hemisphere) would also mediate the effects of music on reading, given the proposed role of CTS in auditory parsing and in line with evidence that stronger left-hemispheric processing is associated with enhanced language and reading outcomes^42,43^.

## Results

### Group Differences

To assess baseline differences between the Music and Control groups, we conducted Mann-Whitney *U* tests across several behavioural and neural variables (see *Table 1*). Compared to controls, the Music group demonstrated significantly higher *Reading Scores* (*U* = 587.5, *p* = .017), *PA Scores* (*U* = 568.5, *p* = .030) and *Musicality Scores* (*U* = 806.0, *p* < .0001). No significant differences were found between groups in *Age, Gender, SES*, *WM Scores*, *Executive Function*, *Tapping Variability, Left*– and *Right-Hemispheric CTS or Left-Lateralised CTS*.

**Table 1.** Descriptives and group differences in demographics, questionnaire, behavioural and neural measures.

|  | Music |  | Control |  |  |  | FDR |
| --- | --- | --- | --- | --- | --- | --- | --- |
|  | *M* | *SD* | *M* | *SD* | *U* | *p* | *p* |
| **Demographics** |  |  |  |  |  |  |  |
| Age | 7.51 | 1.11 | 7.36 | 0.89 | 470.5 | 0.283 | 0.445 |
| Gender | 1.54 | 0.51 | 1.48 | 0.51 | 425.0 | 0.691 | 0.760 |
| **Questionnaires** |  |  |  |  |  |  |  |
| Socio-Economic Status | 0.84 | 0.11 | 0.79 | 0.19 | 421.5 | 0.771 | 0.849 |
| Executive Function | 13.27 | 8.45 | 16.81 | 9.68 | 317.0 | 0.170 | 0.318 |
| Musicality | 2.63 | 0.33 | 0.34 | 0.56 | 806.0 | <0.001** | <0.001** |
| **Behavioural Tasks** |  |  |  |  |  |  |  |
| Reading | 39.12 | 8.13 | 28.74 | 14.37 | 587.5 | 0.003* | 0.017* |
| Phonological Awareness | 44.38 | 10.28 | 34.48 | 15.10 | 568.5 | 0.008* | 0.030* |
| Working Memory | 1.42 | 0.34 | 1.52 | 0.34 | 317.5 | 0.173 | 0.318 |
| Tapping Variability | 0.49 | 0.01 | 0.49 | 0.02 | 458.0 | 0.383 | 0.526 |
| **Neural Variables** |  |  |  |  |  |  |  |
| Left-Hemispheric CTS | 0.00133 | 0.00057 | 0.00135 | 0.00068 | 407.5 | 0.949 | 0.949 |
| Right-Hemispheric CTS | 0.00122 | 0.00065 | 0.00161 | 0.00089 | 295.5 | 0.086 | 0.238 |
| Left-Lateralised CTS | 0.00012 | 0.00072 | -0.00026 | 0.00062 | 525.5 | 0.051† | 0.151 |

*Note.* \* $p < 0.05$ , \*\* $p < 0.001$ . Table includes Mann–Whitney U tests and False Discovery Rate (FDR) corrected $p$ values. CTS = Cortical Tracking of Speech.

Additionally, for the CTS measures, MI’s significance thresholds were determined via permutation testing with 1000 shuffled speech envelopes to help determine whether the observed MI is greater than what would be expected by chance. The average of the latency (or lag) of the MaxMI value for all participants was observed as typical values (for the left hemisphere: *M* = 0.10, *SD* = 0.08; and for the right hemisphere: *M* = 0.10, *SD* = 0.7) for a young adult healthy population as per previous literature^44^.

### Predictors of Reading

To examine the contribution of demographic, behavioural, and neural variables on reading ability, a ridge regression model was conducted to predict reading ability (see *Table 2* and *Figure 1*).

**Figure 1.**
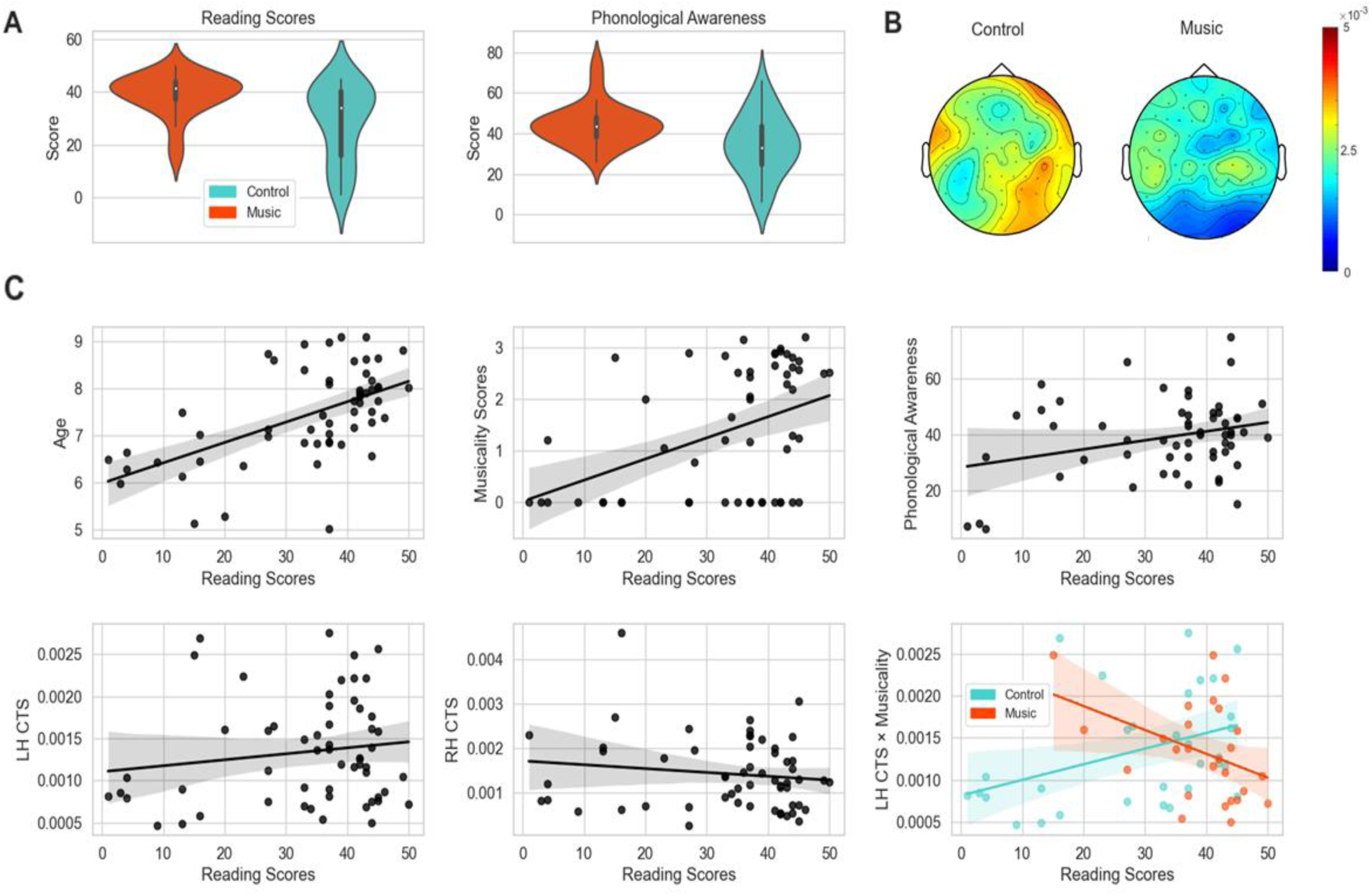
Group differences in reading and phonological awareness, EEG speech processing topographical plots, and predictors of reading. *Note. LH* = Left Hemisphere, *RH* = Right Hemisphere, *CTS* = Cortical Tracking of Speech. **A** Violin plots depicting the distribution (mean and standard deviation) of scores on the Word Identification Fluency task and Phonological Awareness task, across Music (red) and Control (blue) groups. **B** Topographic EEG heatmaps for both Music and Control groups visually reveal generally lower CTS values (measure by Mutual Information magnitude) in the Music group compared to Controls, with neural activity more concentrated over left temporoparietal channels. Control group CTS: LH Min = 0.000467, LH Max = 0.002752; RH Min = 0.000275, RH Max = 0.004580. Music group CTS: LH Min = 0.000496, LH Max = 0.002494; RH Min = 0.001539, Max = 0.002701. **C** Significant predictors of reading scores (in a Word Identification Fluency task) include age, phonological awareness, RH CTS, LH CTS, and the interaction of LH CTS with Musicality scores.

**Table 2.** Summary table for the Ridge Regression model predicting Reading scores.

| Feature | Coefficient | SE | 95% CI |  | *p* | Std.<br>Beta | Partial<br>*R*<sup>2</sup> |
| --- | --- | --- | --- | --- | --- | --- | --- |
|  |  |  | LL | UL |  |  |  |
| Intercept | 0.83 | 0.18 | 0.48 | 1.19 | **<.001 \*\*** |  |  |
| **Demographics** |  |  |  |  |  |  |  |
| Age | 0.77 | 0.28 | 0.19 | 1.34 | **0.011\*** | 0.77 | 0.19 |
| Gender | -0.26 | 0.20 | -0.67 | 0.15 | 0.204 | -0.26 | 0.05 |
| **Questionnaires** |  |  |  |  |  |  |  |
| Socio-Economic Status | -0.06 | 0.19 | -0.44 | 0.32 | 0.761 | -0.06 | 0.00 |
| Executive Function | -0.03 | 0.21 | -0.46 | 0.40 | 0.878 | -0.03 | 0.00 |
| Musicality Score | 1.05 | 0.50 | 0.04 | 2.06 | **0.042\*** | 1.05 | 0.12 |
| **Behavioural Tasks** |  |  |  |  |  |  |  |
| Working Memory | 0.09 | 0.24 | -0.40 | 0.57 | 0.713 | 0.09 | 0.00 |
| Tapping Variability | -0.13 | 0.22 | -0.57 | 0.31 | 0.546 | -0.13 | 0.01 |
| Phonological Awareness | 0.41 | 0.18 | 0.04 | 0.78 | **0.029\*** | 0.41 | 0.14 |
| **Neural Variables** |  |  |  |  |  |  |  |
| Left-Hemispheric CTS | 1.41 | 0.35 | 0.70 | 2.12 | **<0.001\*\*** | 1.41 | 0.34 |
| LH CTS × Musicality | -1.56 | 0.53 | -2.64 | -0.49 | **0.006\*** | -1.56 | 0.22 |
| Right-Hemispheric CTS | -0.96 | 0.31 | -1.59 | -0.32 | **0.004\*** | -0.96 | 0.23 |
| RH CTS × Musicality | 0.78 | 0.42 | -0.06 | 1.63 | 0.068 | 0.78 | 0.10 |

*Note.* \**p* < .05, \*\**p* < .001. LH = Left Hemisphere, RH = Right Hemisphere, CTS = Cortical Tracking of Speech. The model includes demographic, questionnaire, behavioural and neural predictors of reading performance in a Word Identification Fluency task.

The regression model included *Age, Gender, SES, WM, PA*, *Executive Function*, *Tapping Variability*, *Left* and *Right-hemispheric CTS*, and their interaction with *Musicality* as predictors for the outcome variable *Reading* scores. Predictor variables were standardized prior to analysis, and the dataset was split into training (80%) and test (20%) sets. To address multicollinearity and improve generalizability, ridge regression with L2 regularization was applied, using a penalty term of *α* = 0.159986) (*Supplementary Information – Figure 1)*.

The model explained a modest but significant proportion of the variance in reading (*R²* = 0.327, *MSE* = 0.806). Among the significant predictors, *PA* (β = 0.41, *p* = .029), *Left-Hemispheric CTS (LH CTS)* (β = 1.41, *p* < .001), its interaction with *Musicality* scores (β = –1.56, *p* = .006), and *Right-Hemispheric CTS* (RH CTS) (β = –0.96, *p* = .004), were all significant predictors of reading scores. *Age* (β = 0.77, *p* = .011) and *Musicality* Score (β = 1.05, *p* = .042) were also positively associated with reading scores. *Gender* (β = –0.26, p = .204)*, SES* (β = –0.06, p = .761)*, WM* (β = 0.09, p = .713)*, Executive Function* (β = –0.03, p = .878), *Tapping Variability* (β = –0.13, p = .546), and the *RH CTS* interaction with *Musicality* (β = 0.78, p = .068) were not significant predictors.

To confirm the robustness of the results and validate our findings’ consistency against previously published research^44^, an additional regression using the mean MI across all electrodes in the left and right hemispheres (rather than maximum of MI across electrodes) was conducted (see *Supplementary Information – Table 1*). This analysis provided similar pattern of results with significant effects observed for LH CTS, RH CTS and the interaction between Musicality and LH CTS, supporting the consistency of the neural predictors across both measures of MI. However, the current model using maximum (peak) MI values showed stronger effect sizes and greater partial R^2^ values for the key neural predictors, indicating that this metric may better capture individual variability in neural tracking of speech in predicting our dependent variable *Reading*.

### Mediation Analysis

Mediation analyses were conducted to examine the pathways through which musicality influences reading ability, controlling for *Age, Gender, SES, WM, and Executive Function*. Results based on 20,000 bootstrapped samples showed a significant total effect of *Musicality* scores on *Reading* (β = 0.504, SE = 0.144, 95% CI [0.215, 0.794], *p* < 0.001) and a significant direct effect (β = 0.301, SE = 0.142, 95% CI [0.015, 0.586], *p* = 0.039), indicating partial mediation (see *Figure 2*).

**Figure 2.**
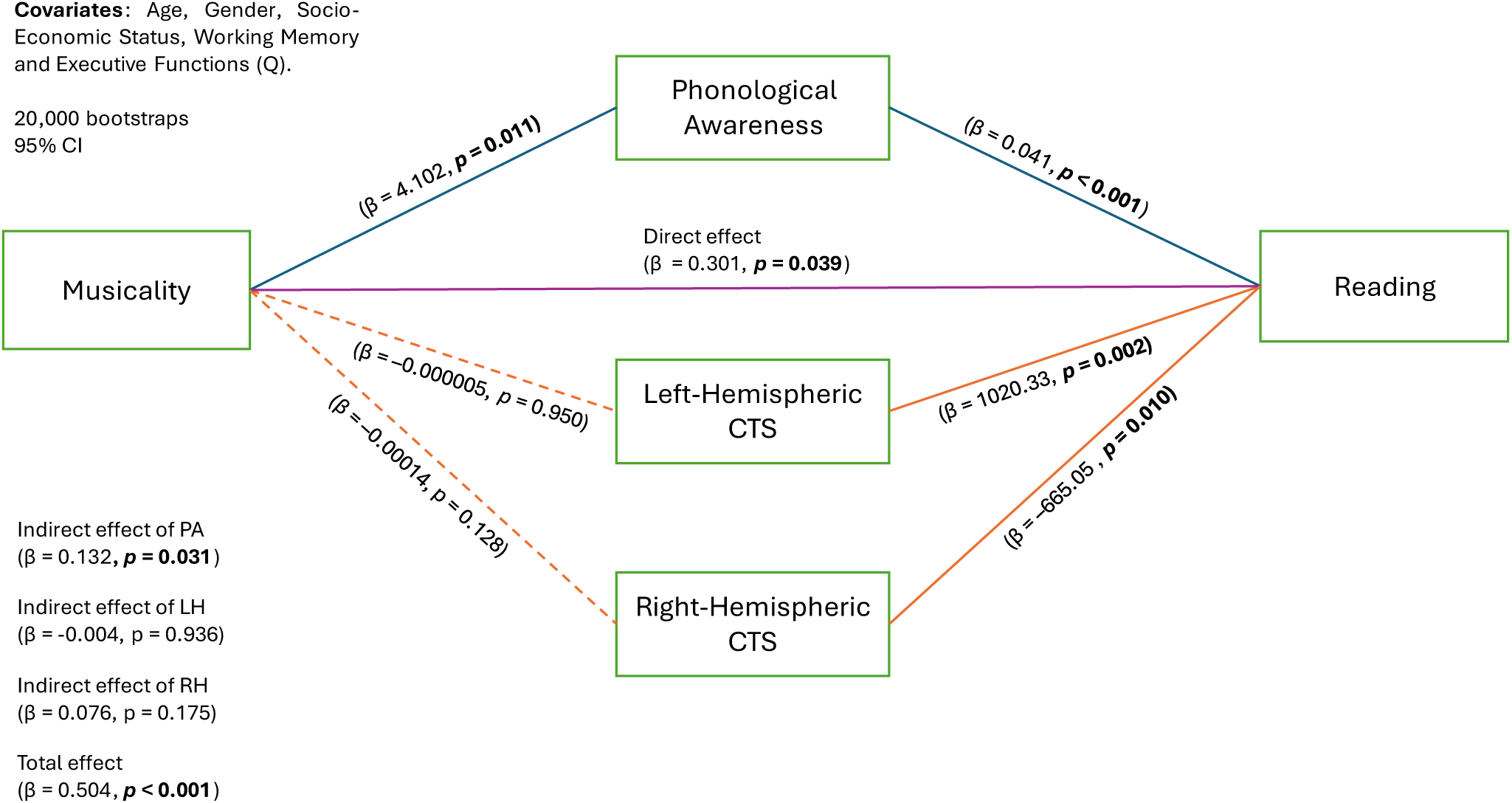
Mediation Pathways linking Musicality to Reading through Phonological Awareness, Left– and Right-Hemispheric Cortical Tracking of Speech. *Note.* Results indicating dependent variable (Reading), independent variable (Musicality) and mediators (PA, Left– and Right-Hemispheric CTS. Straight lines indicate significant effects whereas dashed lines indicate non-significant effects between variables.

*Musicality* significantly predicted *PA* (β = 4.102, SE = 1.546, 95% CI [1.00, 7.21], *p* = 0.011), suggesting children with higher musical training scores performed higher on the *PA* task. In contrast, *Musicality* did not significantly predict *Left* (β = –0.000005, SE = 0.000073, 95% CI [–0.00015, 0.00014], *p* = 0.950) or *Right-Hemispheric CTS* (β = –0.00014, SE = 0.00009, 95% CI [–0.00032, 0.00004], *p* = 0.128).

*Reading* scores were significantly predicted by *PA* (β = 0.041, SE = 0.012, 95% CI [0.018, 0.064], *p* < 0.001) as well as by both *Left-Hemispheric CTS* (β = 1020.33, SE = 313.93, 95% CI [389.13, 1651.52], *p* = 0.002) and *Right-Hemispheric CTS* (β = –665.05, SE = 247.61, 95% CI [–1162.90, –167.20], *p* = 0.010).

The indirect effect of *Musicality* on *Reading* was significantly mediated by *PA* (β = 0.132, SE = 0.083, 95% CI [0.008, 0.334], *p* = 0.031), whereas the indirect effects through *Left*-(β = – 0.004, SE = 0.070, 95% CI [–0.150, 0.138], *p* = 0.936) and *Right-Hemispheric CTS* (β = 0.076, SE = 0.065, 95% CI [–0.007, 0.266], *p* = 0.175) were not significant. These results suggest that musicality impacts reading ability primarily through *PA*, with *CTS* contributing less directly.

## Discussion

The present study examined the impact of musical training on early reading development and its underlying neurocognitive mechanisms, with a focus on phonological awareness (PA) and cortical tracking of speech (CTS). To our knowledge, this is the first study to jointly examine musical training, cortical tracking of continuous naturalistic speech, PA, and reading outcomes in children who underwent musical training, providing a developmentally grounded window into the neural mechanisms linking music and literacy. Four key findings emerged. First, musically trained children outperformed controls on PA and Word Identification Fluency tasks, supporting our first hypothesis and consistent with theories linking music and language development^1,13,14^. Second, unlike our initial hypothesis, CTS did not significantly differ between the musically trained and control groups, with a clear trend for Controls to show stronger CTS in both hemispheres (see Figure 1c). Third, individual differences in both left, and right-hemispheric CTS significantly predicted reading outcomes across all participants even after accounting for other behavioural and demographic factors relevant for reading development. Left CTS emerged as a predictor of enhanced reading skills in the Control (not musically trained) group, with the Music group showing higher reading scores at lower levels of left-hemispheric CTS. Fourth, mediation analyses revealed that PA mediated the relationship between musical training and reading suggesting that musical training boosts literacy by increasing sensitivity to phonological structures within the language.

Taken together, these findings provide new evidence for music-to-language transfer frameworks such as the OPERA hypothesis^16^ and the PRISM model^17^, testing both behavioural and neural pathways through which musical training may enhance reading development during childhood. While recent work has questioned far-transfer effects of musical training^25^, our results suggest that musical training, at least in young children, is associated with improved PA, which in turn contributes to reading ability — pointing to a specific developmental pathway linking music and literacy.

### Musical Training Enhances Reading and Phonological Awareness in Young Children

Children in the musical training group outperformed controls on standardized measures of reading and PA, consistent with a growing body of research showing that musical training can support core language and literacy skills^13,14,20,45,46^. These group differences remained significant despite groups being matched on age, gender, SES, and executive function, indicating a specific association between musical experience and language-related outcomes. Notably, these differences emerged in a relatively young cohort (5–9-year-olds)—a critical^34^ period during which phonological skills and foundational reading abilities are rapidly consolidating^47^. While most studies on music-to-literacy transfer have focused on older children or adolescents, few have examined formal musical training during this early window of literacy development, making the current study a distinct contribution to this emerging area.

To test whether musical training promotes auditory-motor integration which has been linked to reading development^39^ we measured metronome tapping synchronisation. The absence of reliable differences in tapping performance between the music and control groups is noteworthy. Importantly, all control participants engaged in extracurricular activities to a similar extent as the music group, and all of them performed sensorimotor activities—extracurricular sports—that are known to support the development of higher-order cognitive functions^48^ and motor timing precision^49^, which could have improved their motor synchronization to match that of the music group. However, this makes the persistence of music-related benefits in reading and PA even more striking. Rather than suggesting a lack of benefits from musical training to motor synchronisation, these findings emphasize the need to distinguish between different forms of early enrichment when interpreting cognitive outcomes. Specifically, while both groups may have gained temporal coordination skills from their respective activities, our results suggest that musical training exerts effects on language-related outcomes that extend beyond those of general sensorimotor or cognitive engagement^50^.

Our key findings emerged from the regression analysis. The strongest behavioural predictors of higher reading scores were age, musicality, and PA. The contributions of age and PA to reading performance were expected, consistent with a large body of prior research^7,18,19,34^. Musicality emerging as a key predictor of reading is consistent with the body of work showing that musical expertise enhances the temporal and spectral resolution of auditory processing, which in turn facilitates more efficient encoding of speech rhythms and phonemic structure^14,16,39^ . These benefits may sharpen the mapping between auditory input and linguistic representations, providing a neurocognitive scaffold for literacy acquisition^1,51^.

### Musicality and Cortical Tracking of Speech Predict Reading Outcomes

As CTS has been proposed to support auditory parsing that positively contributes to the development of phonological skills over time^52^, we expected stronger CTS in younger readers would (a) predict more advanced reading skills and (b) mediate the relationship between musicality and reading. While both right and left-hemispheric CTS and its interaction with musicality predicted reading scores, these effects revealed a more nuanced neural profile underlying literacy development. Specifically, stronger left-hemispheric CTS was positively associated with overall reading performance, in line with models emphasizing the critical role of left-lateralized auditory processing in supporting phonological and reading skills^27,34^. However, this effect was moderated by musicality: among children with higher musical scores, the positive association between left-hemispheric CTS and reading was less pronounced. We interpret this interaction by proposing that musical experience facilitates development of the left hemispheric language-specialised auditory networks in such a way that musically trained children (a) show a trend for stronger shift to left-lateralised CTS (b) involve weaker engagement of the left-lateralised networks performing CTS to achieve the same reading performance as children not engaged in musical training. In other words, musical training may accelerate or stabilize the maturation of left-lateralised speech-tracking mechanisms, thereby reducing the extent to which left-hemispheric CTS individually predicts reading outcomes^43,53^. This interpretation resonates with recent large-scale replication work showing no reliable enhancement of early auditory encoding in musically trained adult participants^33^ and is also supported by broader group-level patterns in our data.

Contrary to our expectations, musically trained children did not exhibit significantly greater CTS than controls; rather, the control group showed a trend toward stronger right-hemispheric CTS. Increased cortical tracking in the right hemisphere, particularly at the acoustic-temporal level, has been linked to compensatory processing or increased listening effort rather than enhanced linguistic proficiency^54^. From a developmental standpoint, this aligns with evidence that CTS strength can decline with increasing language proficiency and neural maturation, reflecting a shift from reliance on low-level, bottom-up acoustic cues to more efficient, top-down predictive strategies in childhood^55^. In this context, the lack of elevated left or right-hemispheric CTS in the musically trained group may reflect more mature or specialised auditory processing mechanisms.

Finally, right-hemispheric CTS negatively predicted reading performance is in line with the work suggesting that efficient literacy development depends on a shift away from right-dominant auditory tracking. This aligns with longstanding neurodevelopmental models in which increasing reading proficiency is associated with greater left-hemispheric specialisation for speech and phonology^27,42^. This interpretation aligns with models of experience-dependent neural tuning, which propose that enriched auditory environments such as music accelerate the specialisation of speech-relevant neural circuits^56,57^.

### Phonological Awareness but not CTS Mediates the Relationship between Musical Training and Reading

PA significantly mediated the relationship between musicality and reading, supporting models positing that musical training enhances reading ability indirectly by sharpening phonological processing skills^13,41,45,58^. Notably, both the indirect path via PA and the total effect of musicality on reading were significant, and the direct path remained significant as well, indicating partial mediation. Together, these findings reinforce the role of PA as a critical link between enriched auditory experience—such as musical training—and early cognitive processes that underpin literacy acquisition^47^. In contrast, neither left nor right-hemispheric CTS mediated the effect of musicality on reading, indicating that while CTS emerged as a predictor of reading in regression models, it does not function as a neural mechanism directly linking musicality to reading outcomes in young readers^33^. One interpretation is that musicality simultaneously promotes left-lateralised CTS while also enhancing reading through phonological skills, with this CTS profile representing a consequence rather than a precursor of reading development (see Destoky et al., 2022 for similar views in dyslexia).

These findings have translational value for early years educational practices. Most children begin formal reading instruction around age 4–6, typically using phonics-based approaches that teach them to map individual speech sounds (phonemes) onto corresponding letter groups (graphemes), which are then blended to form words. This phoneme–grapheme mapping is a foundational skill for literacy acquisition in the early school years and a key predictor of early reading achievement^58,59^. Previous studies in older children have shown that musical training strengthens rhythmic discrimination abilities, which in turn support the development of PA by reinforcing sensitivity to phonemes—ultimately facilitating more efficient decoding and reading skills^13,41^. Our findings show that musicality boosts reading via phonological skills in younger readers (primary school yeas 1-3) as well, independently of other critical behavioural and demographic factors. Yet relatively few children begin structured musical training at the age of 4-6. Interventions targeting this developmental window, when neural systems supporting auditory and phonological processing are especially malleable^4^, may therefore yield stronger and longer-lasting effects than later interventions.

### Future Research

Future studies should aim to explore CTS longitudinally across early development starting before formal reading instruction begins. Musical training longitudinally facilitates greater neural plasticity and flexibility within auditory networks in young adults^24^. Therefore, longitudinal and interventional studies comparing musical training and control groups as early as age 4-6 could clarify whether earlier engagement with musical training produces more robust gains in reading through neuro-cognitive changes in auditory processing. Research should also compare musical training with other high-skill activities (e.g., dance, sports, or theatre) to disentangle domain-specific from domain-general cognitive benefits. Evidence from intervention studies indicates that musical training can enhance PA and improve reading outcomes in both typically developing children and those with reading difficulties^12,60,61^. Including children with dyslexia in future work could clarify whether musical training helps remediate deficits in the temporal sampling of speech—a known neural hallmark of dyslexia— and whether CTS can serve as a sensitive index of such remediation^62^.

### Conclusion

This study provides novel evidence that musical training is associated with enhanced reading, phonological skills as well more left-lateralised cortical tracking of speech in young children at the onset of literacy. Phonological awareness emerged as a key mediator between musicality and reading supporting the hypothesis that musical training boosts language-specific cognitive functions relevant for phonological development. Musicality and left hemispheric cortical tracking of speech further predicted higher reading outcomes. None of the neural measures of cortical speech tracking emerged as mediators between musicality and reading, suggesting that neural mechanisms of auditory speech analysis and parsing either develop in parallel to or as a consequence of reading acquisition. These findings contribute to music-to-language transfer models by demonstrating that musical training sharpens phonological processing in ways that facilitate reading—particularly during a sensitive developmental window when phonological awareness is key for successful reading acquisition and the underlying neural systems are highly malleable. As such, structured musical training may offer a practical, engaging, and low-cost intervention for supporting foundational literacy skills, with particular promise for supporting children at risk for phonological difficulties.

## Methods

### Participants

Sixty-two children participated in the study. Participants were recruited through advertisements in local libraries, social media networks, and the University of Aberdeen’s staff network. No participants reported any visual or auditory impairment or had previously or currently been diagnosed with any neurodevelopmental disorder (e.g., ADHD, Dyslexia, ASD). All participants were residents in the United Kingdom and spoke English as a first language.

Participants who did not fully complete any of the EEG assessments and/or did not meet the minimum requirements for inclusion in the behavioural tasks were excluded from the sample (*N=*5). The final sample consisted of 57 participants (*M*=7.42 years, *SD*=0.99, 28 female), that were divided into two groups: 26 children were assigned to the music group (*M*=7.51 years, *SD*=1.11, 12 female) and 31 children to the control group (*M*=7.36 years, *SD*=0.89, 16 female). The music group included participants involved in extracurricular music activities, such as learning to play an instrument (*N*=23) or taking singing lessons (*N*=3). To ensure both groups had similar extracurricular commitments, the control group participants were also engaged in non-musical extracurricular activities like playing sports, joining scouts, or participating in after-school clubs. All participants were required to have been enrolled in their respective activities for at least one month, with the activities occurring for at least thirty minutes to one hour per week. Participants were matched for age (*U* = 517.5, *p* = 0.261), gender (*U* = 468.0, *p* = 0.712), Caregiver’s level of education (*U* = 458.5, *p* = 0.712), Household Income (*U* = 505.0, *p* = 0.515) and Executive Function (*U* = 326.5, *p* = 0.439). For a more details on the demographic information of participants Table 3.

**Table 3.** Demographic information for participants on each group.

| **Sample** | **Music**<br>***M (SD)*** | **Control**<br>***M (SD)*** |
| --- | --- | --- |
| N | 26 | 31 |
| **Age (Years)** | 7.51 (1.11) | 7.36 (0.89) |
| **Gender** | 1.54 (0.51) | 1.48 (0.51) |
| (1) Female (total) | 12 (28) | 16 (28) |
| **Ethnicity (%)** | 1.23 (0.43) | 1.22 (0.67) |
| (1) White | 77% | 87% |
| (2) Mixed/Multiple groups | 23% | 7% |
| (3) Asian/Asian British | 0% | 3% |
| (4) Other ethnicity groups | 0% | 3% |
| **Caregiver's level of education (%)** | 5.42 (0.86) | 5.38 (0.83) |
| (1) Did not complete any school qualification | 0% | 0% |
| (2) Completed first school qualification at about 16 years (e.g. GCSE/Junior High School) | 0% | 0% |
| (3) Completed post-16 vocational course | 4% | 6% |
| (4) Completed Second qualification (e.g. A levels/ High School) | 0% | 3% |
| (5) Undergraduate degree or professional qualification | 46% | 35% |
| (6) Postgraduate degree | 50% | 55% |
| (0) Prefer not to say | 0% | 0% |
| **Household Income** | 6.54 (2.06) | 6.00 (1.99) |
| (1) Under £20k | 0% | 6% |
| (2) £20-30k | 0% | 3% |
| (3) £30-40k | 4% | 3% |
| (4) £40-50k | 8% | 6% |
| (5) £50-60k | 11% | 3% |
| (6) £60-80k | 19% | 30% |
| (7) £80-100k | 23% | 26% |
| (8) Over £100k | 35% | 23% |
| (0) Prefer not to say | 0% | 0% |
| **Executive Functioning (BRIEF-2)** | 13.27 (8.44) | 16.81 (9.67) |
| Low | 70% | 52% |
| Medium | 26% | 42% |
| High | 4% | 6% |
| **Musicality Questionnaire** | 2.63 (0.33) | 0.34 (0.56) |
| Low | 0% | 71% |
| Medium | 0% | 29% |
| High | 100% | 0% |
| **Hours/week on respective activity** | 1.25 (0.53) | 2.30 (1.13) |

*Note.* For group differences statistics please see Table 1.

Written informed consent was obtained from caregivers of all participants before taking part in the study, both online and in person. Families received a £16 time reimbursement, a sticker, and a pencil for participating. The study was approved by the University of Aberdeen Ethics Committee (approval code 1271479) and was conducted in accordance with the principles outlined in the Declaration of Helsinki. Researchers involved in the data collection process underwent checks from the Protecting Vulnerable Groups (PVG) scheme managed by Disclosure Scotland.

### General Procedure and Apparatus

Before coming to our laboratory, caregivers completed online questionnaires related to their household and child, which took approximately 30 minutes to complete. Children then participated in an in-person session where they completed behavioural and neurophysiological tasks. The session lasted for 1 hour including instructions and break time. Caregivers were asked to not intervene when tasks took place and not to aid children in completing the tasks. Children sat on a desk next to an experimenter while caregivers waited outside. Auditory stimuli were presented over loudspeakers (with same volume settings across participants). A computer keyboard was provided to children to respond to stimuli when required and for the experimenter to compute responses and move between tasks. Audio was recorded during reading and phonological tasks for posterior check of responses. Children took short breaks in between tasks if needed. The administration of tasks ensured a standardized approach, maintaining consistency in assessment across all participants.

The online platform Testable^63^ (www.testable.org) was used to acquire information from parent-reported questionnaires. Parents were asked to find a comfortable and quiet space and to allocate the recommended time to complete the questions from a personal electronic device connected to a web server. Psychopy2 Builder software^64^ was utilized to program and administer the tasks during the test sessions. During test sessions, stimuli was presented using a computer screen in a noise-cancelling booth. EEG recordings took place in a sound-attenuated, noise and signal-cancelling booth. EEG data were recorded from 64 scalp electrodes and 8 external electrodes, two of which were positioned to monitor cardiac activity, using the Active Two system (Biosemi, Amsterdam, Netherlands). Scalp electrode placement followed the international 10/20 system. EEG processing and cleaning was carried out using MNE Python (version 1.6)^65^. Auditory envelope extraction and Mutual Information analyses were conducted using MATLAB (2023. Version R2023a. The MathWorks Inc., Natick, MA, USA).

### Questionnaires

#### Demographics

Demographic questions were acquired from Politimou et al., (2018)^66^. The demographics questionnaire included questions related to caregivers, children, and their household such as languages spoken at home, parents’ education, household income, among others.

Factors related to Socio-Economic Status (SES) play a significant role during development, as higher SES is associated with greater access to music education and, consequently, better academic outcomes ^2,67^. To assess SES, we derived two main indicators: caregiver’s education scores and income scores. Parent education scores were based on the highest level of education attained by the primary caregiver, ranging from a minimum score of 1 (indicating no formal qualifications) to a maximum score of 6 (representing a postgraduate degree). Income scores ranged from 1 to 8, with a score of 1 indicating an annual income of less than £20,000 and a score of 8 representing an annual income exceeding £100,000. Both scores were rescaled to a common 0–1 range using min-max normalization, according to the formula: *Xscaled = (X – Xmin) / (Xmax – Xmin).* The final SES score was calculated by averaging the rescaled education and income scores, resulting in a composite SES measure in which higher values represent higher socioeconomic status. Higher scores indicated higher SES.

#### Musical Training and Engagement

Children in both the music group and non-music control group (those that either played an instrument, enrolled in singing or had ballet classes regularly) completed a questionnaire designed to assess their degree of musical training and involvement. The questionnaire included information on musical activities, inclusive of playing an instrument, practice time, length of time playing, confidence playing, and engagement in other extracurricular musical activities such as concerts or recitals.

Responses were given in a Likert style scale, questions rating from 1 to 10. Scores were calculated from numerical responses with a higher score indicating greater musical training and engagement. To calculate the sum of all measures, we normalized each question, scaling them to a range of 0 to 1 using a min-max scaling approach.

#### Executive Functions

Musical training may enhance literacy through multiple pathways, including domain-general executive functions and language-specific processes. Learning to play an instrument during childhood engages Executive Function skills such as working memory and attentional control^50,68^ (for a review see Rodriguez-Gomez & Talero-Gutierrez, 2022), which are known to predict reading skills^69^ and support reading comprehension^70^. To assess difficulties in these domain-general cognitive skills, children’s executive abilities were evaluated using the Behavior Rating Inventory of Executive Function (BRIEF2)^71^. This parent-report questionnaire measures executive dysfunction and includes subscales assessing key executive function components such as self-monitoring, inhibition, shifting, emotional control, initiation, working memory, planning/organization, task-monitoring, and organization of materials.

Questions were rated in a Likert scale from “Never a problem”, “Sometimes a problem” and “Often a problem” (rated from 0 to 2 respectively). Scores from all questions were summed giving a total Executive Function score. A higher score on the BRIEF2 indicates poorer executive function abilities. Higher scores in this test indicate some degree of executive dysfunction.

### Tasks and Stimuli

#### Phonological Awareness

A phonological awareness task was administered to assess participants’ proficiency in identifying and articulating phonemes within words. The task aimed to evaluate the participants’ sensitivity to the sound structure of language, specifically focusing on their ability to segment and manipulate individual sounds within words. Words were acquired from DIBELS, University of Oregon^72^, (see also *Supplementary Materials Methods 1*) and ranged from 2 to 4 phonemes maximum with a maximum of 24 words and 146 phonemes.

Children were presented with a word at a time and were instructed to pronounce as many correct phonemes per word as possible within a 1-minute period. The experimenter first read the word on the screen and waited for 3 seconds for a response. If the child did not start saying a phoneme in the first 3 seconds, the experimenter will move on to the next word. Answers were coded from audio recordings. Each correct phoneme was recorded as 1 point, otherwise as 0. The PA measure was the sum of all phonemes identified correctly from all words within 1 minute. A high score indicated higher phonological ability.

#### Reading Task

Reading proficiency in children was assessed using a Word Identification Fluency Task^73^. This task is designed to measure the participants’ ability to recognize and pronounce words efficiently. In this task, children were asked to read aloud as many words as they could within a 1-minute period. The words used were selected from the Dolch word list^74^ (https://sightwords.com/sight-words/dolch/#lists), which is segmented into different lists corresponding to the educational grade levels of the children. The list combined words from grades below (e.g. Grade 1), above (e.g. Grade 3) and current educational grade (e.g. Grade 2) of the child at the time of test and words were assigned a frequency of appearance in accordance with their difficulty. To see the words used for each grade, see *Supplementary Materials Methods 2*.

Children were required to read words one by one on the screen while experimenters recorded correct and incorrect answers using a computer keyboard. Answers were coded from audio recordings. Each correct pronouncement of the word was coded as 1, otherwise 0. A high score indicated higher reading proficiency.

#### Working Memory Task

To account for these broader cognitive influences on reading development, we included a visuo-spatial working memory task to behaviourally assess domain-general executive function. The visuo-spatial WM maintenance task followed the protocol used by Sato et al. (2018)^75^.

Participants were presented with two coloured squares (sample stimulus) for 250ms, followed by a fixation cross for 1000ms (retention period), where they were required to remember the previous colours in the squares. After this, two coloured squares will reappear (same colour to previous ones or different) with a question mark as fixation point (test stimulus). During the test stimulus trials, participants will indicate with a computer keyboard whether the sample and test stimuli were the same or different. Participants will press the Tick key, when they matched and Cross key when they did not (keys m and n were covered with stickers). Participants were instructed to respond with their dominant hand using the index and middle fingers. The test stimulus period lasted for 2000ms or until a response was given. After the test stimulus, a fixation cross appeared as an inter-stimulus interval (ISI) that could last between 1250ms±150ms.

There was a total of 100 experimental blocks, 50 matched blocks (row A in Figure 3) and 50 unmatched blocks (row B in *Figure 3*) and the total duration of the task was 7 minutes. At the beginning of the task, the experimenter showed an image like Figure 2 and instructed the participant to press the correspondent keys during test stimulus for matched and unmatched trials. Before starting the experimental blocks, children completed a training phase where they practiced and familiarized themselves with the procedure. The training phase included 10 blocks of 5 matched and 5 unmatched trials.

**Figure 3.**
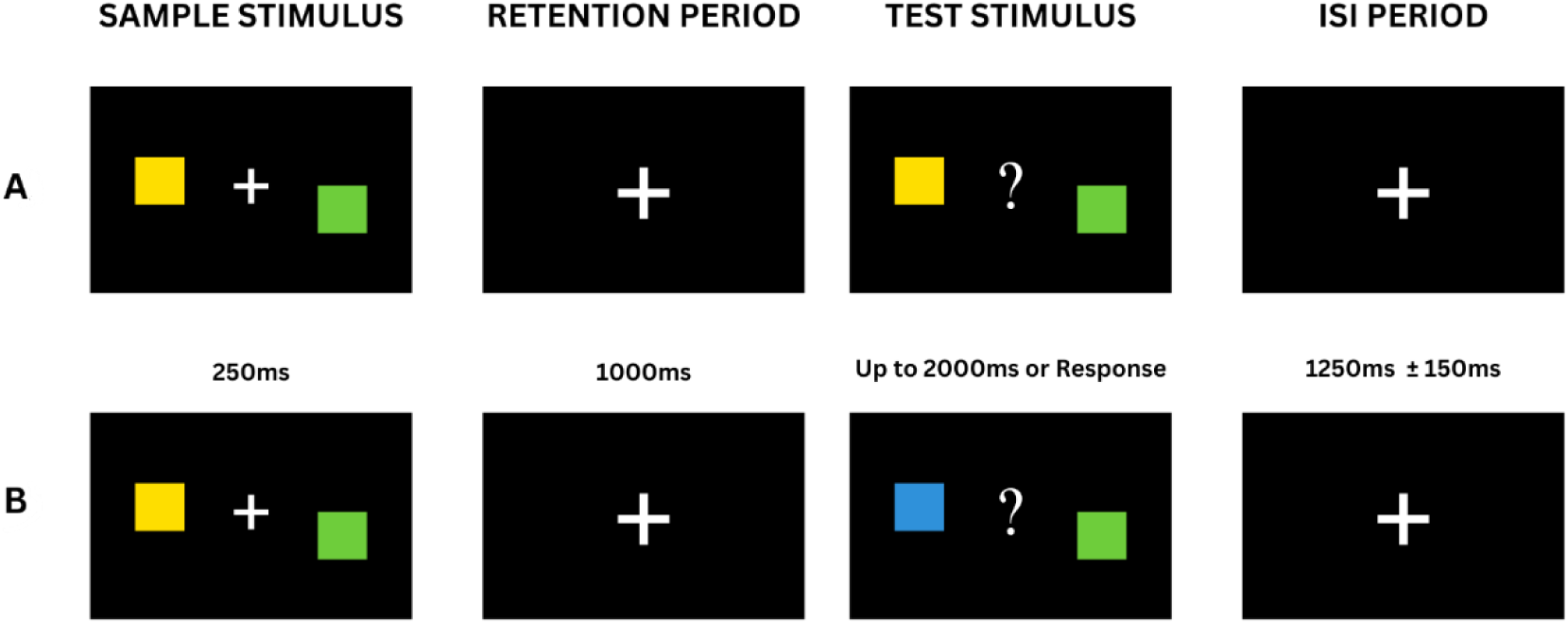
Stimuli sequence for the working memory maintenance task or match-to-sample task from Sato et al. (2018)^75^. *Note*. **A** Example of matching blocks (congruent). **B** Example of unmatching blocks (incongruent).

We focused on calculating accuracy and reaction times (RT) during the test stimulus phase. To assess each participant’s performance in the task, accuracy was calculated as the percentage of correct responses out of the total number of trials (both matched and unmatched blocks) and RT were measured for correct trials only. Children were required to perform a minimum of 50% accuracy to be included in the analysis. An inverse efficiency score (IES) was computed by dividing mean RT by accuracy, such that lower scores indicate more efficient performance. This IES was used as the overall variable referred to as *WM Score*.

#### Tapping Task

This task combines elements from Kalashnikova et al. (2021)^21^, and the Expressive Rhythm: Metronome task by Thomson and Goswami (2008)^76^. There were two types of trials: *Paced* and *Unpaced*.

*Paced trials. T*he auditory stimuli used was a 20-second sequence of pure tone events, each lasting 10 milliseconds and occurring at a rate of 2Hz (500ms). *Unpaced trials.* Each *Paced* trial was followed by a 20-second sequence without auditory stimulus, during which participants continued tapping at the same rate as in the *Paced* trials. A fixation cross appeared on the screen during the trial. After each *unpaced* trial, a feedback screen displayed the message: “*Well done!.”* This block of *paced* and *unpaced* trials was repeated twice. Each block started with a countdown to prepare participants for their response (see *Figure 4)*.

**Figure 4.**
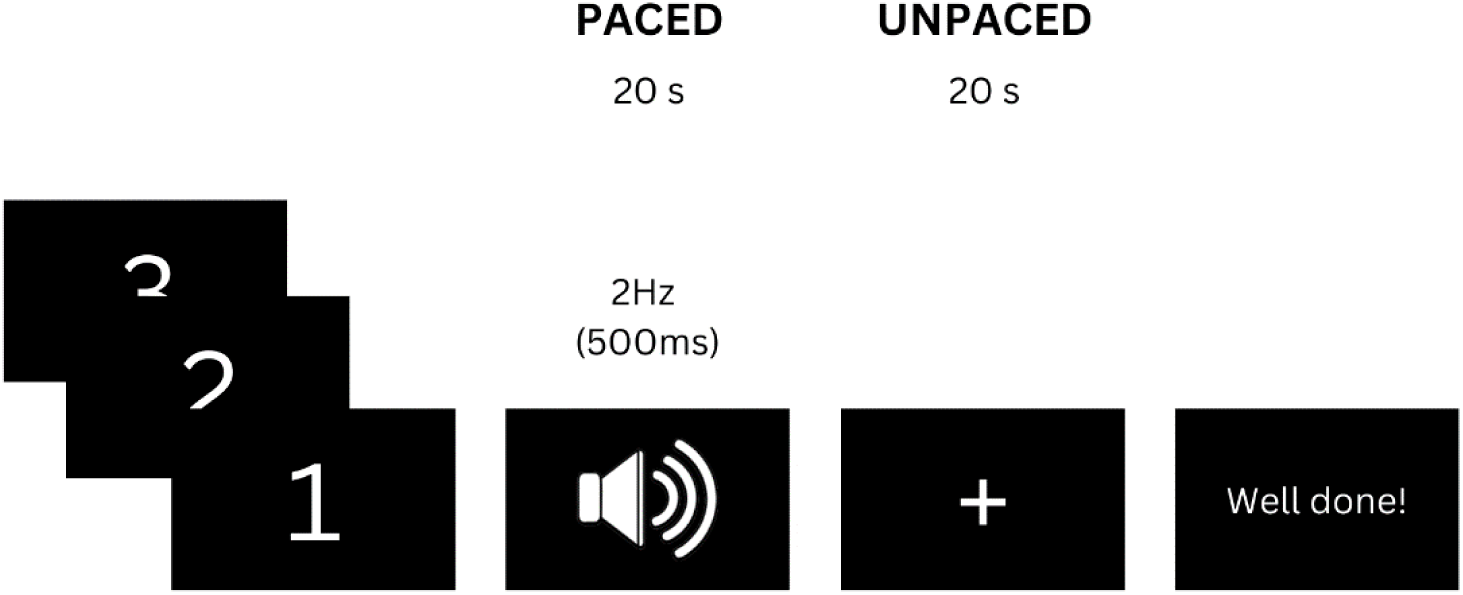
Visual representation of a block from the tapping task.

Participants were instructed to tap along with a metronome beat for a specific duration, aiming to synchronise their taps as closely as possible to the metronome’s tempo. First, participants completed a *familiarization phase* where the experimenter explained the task and its procedures. The experimenter emphasized that they had to tap *in sync* with the sound and to maintain the same rhythm even after the sound stopped, continuing until the cross on the screen disappeared. The experimenter mimicked the sound and tapping as in the experimental trials, and children were asked to tap along with their index finger on the desk. When participants were comfortable and understood the procedure, we proceeded with the *practice phase*, where they completed one block of *paced* and *unpaced* trials while the Space bar on the computer keyboard recorded taps as input by the software. Following this, participants completed two blocks of *paced* (40 s) and *unpaced* trials (40 s).

To ensure the analysis included only taps that were both preceded and followed by metronome events, the first and last taps in the sequence were removed. We first calculated the mean inter-tap interval (ITI) for each participant in both types of trials. These scores were computed by calculating the average time interval in milliseconds between consecutive taps for each participant, following the metronome at a rate of 500 ms, or a tap every 500 ms. Then we calculated the standard deviation (SD) of the ITIs for each participant in both trial types.

For the current study, we only calculated the SD of the ITIs only for *paced trials* which we will refer as *Tapping Variability*. This measure was used as an index of inter-subject variability in performance, capturing the consistency of tapping rates when listening to a metronome beat. A lower *Tapping Variability* indicated greater consistency (i.e., more regular tapping intervals), while a higher value indicated greater inconsistency (i.e., more variable tapping intervals).

### Speech Processing Task

While EEG responses were recorded continuously, children listened to a 5-minute adaptation of *The Gingerbread Man* story (300s; 682 words) read by a male speaker and presented through speakers. To maintain children’s engagement and minimize eye movement artifacts, a story-related image was presented and remained on screen every 30-40s for the duration of the story. The images were designed to encourage to visually focus on the screen while listening to the story. Caregivers accompanied their children if the child requested their presence. They were instructed not to intervene or engage with the child during the task.

#### Envelope Extraction

Auditory envelope extraction, EEG processing and MI analyses were conducted using MATLAB. The auditory signal was first divided into eight frequency bands, spaced equally on a cochlear frequency map between 100 Hz and 8000 Hz, using a filter bank^77^. Each band was bandpass filtered using third-order Butterworth filters and the Hilbert transform was applied to obtain the analytic signal. The envelope for each band was computed as the absolute value of its analytic signal. These narrow-band envelopes were then averaged to produce a single broadband envelope. The resulting envelope was bandpass filtered between 0.5 and 8 Hz to match EEG preprocessing and then resampled to 128 Hz. Finally, the envelope was then normalized using a Gaussian copula transform^78^ and truncated to remove the 500 ms initial offset responses^79^.

#### EEG Processing

The raw EEG data were first loaded, bad channels were rejected based on visual inspection, data was re-referenced to average and bandpass filtered between 0.2 Hz and 40 Hz. Independent component analysis (ICA) was applied to the filtered data (for this step, bandpass-filtered between 1 and 40 Hz to minimise slow drifts and improve components decomposition) to isolate and remove artefactual components related to vertical (blinks) and horizontal eye movements and heartbeat artefacts. Components associated with electrooculography (EOG) artifacts were detected and rejected using automated methods (based on high correlation with the EOG), any additional components were identified and rejected based on visual inspection and correlation maps. ICA was then applied to the original 0.2-40 Hz filtered data. For MI analysis, we used the FieldTrip Toolbox^80^ in MATLAB. EEG data was down-sampled to 128 Hz to match the speech envelope sampling rate and bandpass filtered using third-order Butterworth filters (0.5–8 Hz) to focus on the key frequency contributors of cortex-to-speech envelope tacking. The EEG data were then segmented and aligned with the stimulus, compensating for constant trigger delay (26 ms) between presentation of stimulus and EEG recording times. The Hilbert transform was applied to extract the analytic signal, and the real and imaginary components were normalized using a Gaussian-Copula transformation^78^.

#### Channel Selection

Since temporal and parietal regions are known to be involved in speech processing and have been shown to be most sensitive to speech envelope tracking and MI analysis, we focused the MI analysis on channels over these areas. Specific channels were chosen based on prior research demonstrating their role in cortical speech tracking^44^. For the left hemisphere these included FT7, FC5, FC3, T7, C5, C3, TP7, CP5, CP3, P7, P5, P3, P9, and for the right hemisphere, FT8, FC6, FC4, T8, C6, C4, TP8, CP6, CP4, P8, P6, P4, P10 (see *Figure 5*). This selection ensured that the MI analysis focused on the neural response to speech rather than visual-motor activity.

**Figure 5.**
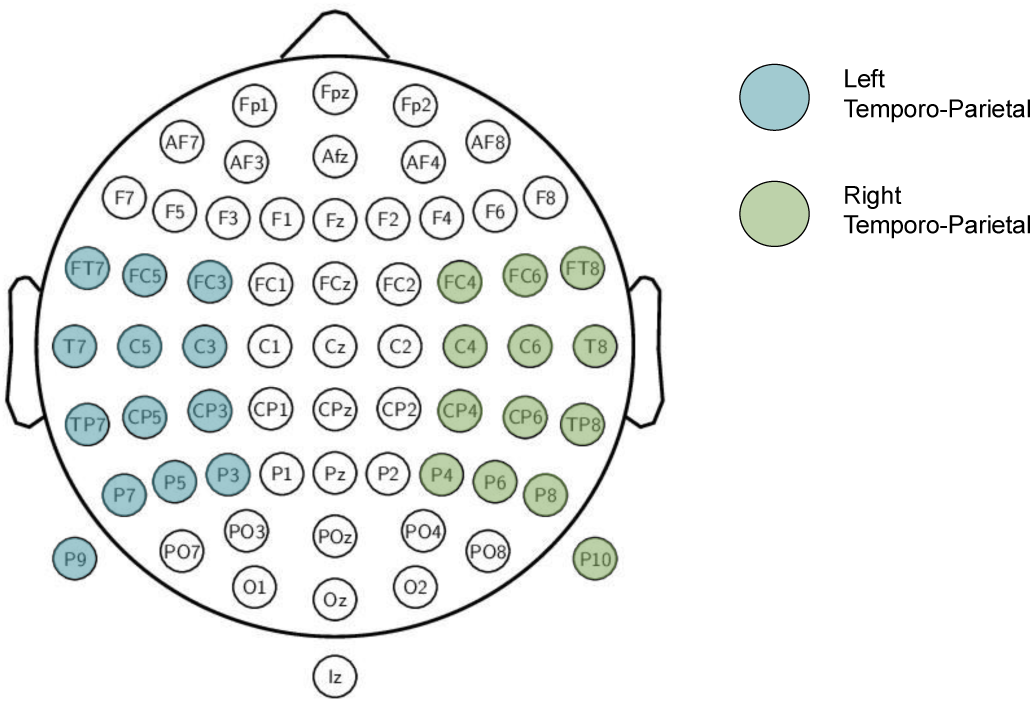
Visual representation of regions of interest and EEG channel selection.

#### MI Analysis

We employed a Mutual Information (MI) approach to assess EEG-based neural tracking of continuous speech. MI techniques have previously been shown to effectively capture nonlinear neural tracking of continuous speech envelopes^44^.

MI between continuous EEG signals and the speech envelope was estimated using the Gaussian copula-based MI method (GCMI)^78,81^. Conceptually, MI is a measure that quantifies how much information one signal contains about another (in this case, how much information the EEG signals contain about the speech envelope) by measuring the reduction in uncertainty about one signal given knowledge of the other. This approach is tightly linked to that of entropy and accounts for linear and non-linear dependencies also being robust to differences in joint and marginal distributions.

MI was computed across a range of time lags (0–200 ms) by shifting the EEG signal relative to the speech envelope in 7.8 ms steps (1 sampling point at 128 Hz). Analyses were conducted separately for different groups of channels corresponding to scalp regions of interest (e.g., left/right temporo-parietal). Within each region, MI was estimated for each participant, channel and lag separately. To summarize these results, we extracted the maximum MI (MaxMI) for each channel. This MaxMI value represents the highest MI observed for that channel across all time lags, indicating the time lag at which the EEG signal aligns most strongly with the speech envelope. Then, we selected the channel with the highest MaxMI in that region, meaning the region-level MaxMI represents the strongest speech-brain coupling observed at any single channel within the region. This measure highlights the peak MI response within each scalp region, rather than an average across channels. We chose this approach because averaging across electrodes can be biased by individual variability in head shape in young children and electrode positioning, potentially reducing sensitivity to the strongest local speech-brain coupling.

MI variables for further analysis included MI analyses separately for the left hemisphere and right hemisphere, which we will refer as to *Left-Hemispheric CTS* (LH CTS) and *Right-Hemispheric CTS (RH CTS)* respectively. Additionally, we calculated how much greater the cortical channels in the left hemisphere track speech in comparison to the right hemisphere, which we will refer to as Left-Lateralised CTS (subtracting left hemisphere MaxMI from the right hemisphere MaxMI values within a participant) as we expect the frontotemporal language network to become increasingly more left-lateralized with age^9^.

### Analysis Plan

Our analysis plan was structured to address the study’s core hypotheses. Before we modelled any relationships between our demographic, questionnaire, behavioural and neural measures, we conducted group differences analyses on all key variables. Due to non-normal distributions, we used Mann–Whitney U tests for the group comparisons and controlled for multiple comparisons using false discovery rate (FDR) correction. These comparisons aimed to determine whether the Music group exhibited enhanced profiles in cognitive, behavioural, and neural measures^1,8,51^, consistent with our first two hypotheses.

To assess the contribution of cognitive and neural variables to reading performance, we employed a regularised linear ridge regression model^82,83^. The model included standardized demographic (*Age*, *Gender*), questionnaire (*SES*, *Musicality*, *Executive Function*), behavioural (*PA, WM, Tapping Variability*), neural predictors (*Left* and *Right-hemispheric CTS*) and their interaction with *Musicality* scores. Reading scores were logit-transformed after min–max scaling to address non-normal distribution, as confirmed by visual inspection of histograms, Q-Q plots, and skewness values. This transformation expands the scale to the full real number line, improving linearity and normality assumptions for regression. Ridge regression was chosen over ordinary least squares due to moderate to high correlations among predictors (see *Supplementary Information – Figure 1* and *Figure 2*), which raised concerns about multicollinearity. By applying L2 regularization, ridge regression penalizes large coefficients, reducing variance and improving model stability, thereby preventing overfitting, and enhancing generalizability. Model assumptions were thoroughly evaluated: the Durbin-Watson statistic indicated no significant autocorrelation of residuals; the Breusch-Pagan test indicated no heteroskedasticity; residuals approximated normality based on Q-Q plots; Variance Inflation Factors were within acceptable ranges for most predictors; and no extreme outliers were observed, supporting the use of ridge regression.

The aim was to determine whether language-specific mechanisms (PA and CTS) could explain the influence of musical engagement on reading ability, consistent with prior literature^13,41,43^. To test whether PA and CTS mediate the effect of musical training on reading, we performed parallel mediation analyses using the *Pingouin* statistical package in Python^84^. This approach allowed us to estimate indirect effects using non-parametric bootstrapping (20,000 resamples), which does not assume the normality of the sampling distribution of indirect paths. The mediation model included *Musicality* scores as the independent variable, *Reading* accuracy as the dependent variable, *PA*, *Left* and *Right-hemispheric CTS* as parallel mediators, and *Age*, *Gender*, *SES, Executive Function* and *WM* scores as covariates.

## Acknowledgements

We sincerely thank all the families, both caregivers and their children, who so generously gave their time and effort to participate in this study. Their engagement made this research possible. We also thank Marta Brzeska and Isla Ramsay for their voluntary support in data collection during their undergraduate studies at the University of Aberdeen. M.C.G.S. and A.K.G. are supported by funding from the School of Psychology, University of Aberdeen and the Research Committee. B.M. is supported by the UKRI Economic and Social Research Council [grant number UKRI628]. A.K. is supported by the Medical Research Council [grant number MR/W02912X/1]. A.K. and A.K.G are members of the Scottish-EU Critical Oscillations Network (SCONe), funded by the Royal Society of Edinburgh (RSE Saltire Facilitation Network Award to A.K., Reference Number 1963).

The authors declare no competing financial interests.

## Author Contributions

M.C.G.S.: Conceptualization, Methodology, Software, Formal Analysis, Investigation, Resources, Data Curation, Writing –Original Draft, Writing –Reviewing & Editing, Visualization, Project Administration and Funding Acquisition. A.K.G.: Conceptualization, Methodology, Software, Investigation, Resources, Writing – Review & Editing, Supervision, Project Administration and Funding Acquisition. B.M.: Conceptualization, Methodology, Writing –Review & Editing, Supervision, Project Administration. A.K.: Software, Writing –Review & Editing, Supervision. All authors reviewed and approved the final manuscript.

## Data Availability

The data supporting the findings of this study are available from the corresponding authors upon reasonable request; see Author Contributions for details on specific datasets and resources.

