## Supplementary Information for "Benefits of music training for learning to read: Evidence from cortical tracking of speech in children"

**Table 1**

*Summary table for the Ridge Regression model predicting Reading scores.*

| Feature | Coefficient | SE | 95% CI | | *p* | Std. Beta | Partial $R^2$ |
|---|---|---|---|---|---|---|---|
| | | | LL | UL | | | |
| Intercept | 0.86 | 0.18 | 0.50 | 1.22 | **<.001 **** | | |
| **Demographics** | | | | | | | |
| Age | 0.84 | 0.29 | 0.26 | 1.43 | **0.006*** | 0.84 | 0.21 |
| Gender | -0.18 | 0.20 | -0.59 | 0.22 | 0.366 | -0.18 | 0.03 |
| **Questionnaires** | | | | | | | |
| Socio-Economic Status | -0.01 | 0.19 | -0.39 | 0.37 | 0.958 | -0.01 | 0.00 |
| Executive Function | -0.03 | 0.21 | -0.47 | 0.40 | 0.882 | -0.03 | 0.00 |
| Musicality Score | 0.95 | 0.49 | -0.03 | 1.94 | 0.058† | 0.95 | 0.11 |
| **Behavioural Tasks** | | | | | | | |
| Working Memory | 0.20 | 0.24 | -0.29 | 0.70 | 0.414 | 0.20 | 0.02 |
| Tapping Variability | -0.12 | 0.22 | -0.57 | 0.32 | 0.572 | -0.12 | 0.01 |
| Phonological Awareness | 0.39 | 0.18 | 0.02 | 0.77 | **0.040*** | 0.39 | 0.13 |
| **Neural Variables** | | | | | | | |
| LH CTS | 1.26 | 0.32 | 0.61 | 1.92 | **<0.001**** | 1.26 | 0.32 |
| LH CTS × Musicality | -1.19 | 0.47 | -2.15 | -0.24 | **0.016*** | -1.19 | 0.17 |
| RH CTS | -0.83 | 0.29 | -1.43 | -0.23 | **0.008*** | -0.83 | 0.20 |
| RH CTS × Musicality | 0.62 | 0.39 | -0.18 | 1.41 | 0.123 | 0.62 | 0.07 |

*Note.* **p* <.05, ***p*<.001. Model performance statistics: $R^2$ Score = 0.1835, Mean Squared Error (MSE) = 0.9768. LH = Left Hemisphere, RH = Right Hemisphere, CTS = Cortical Tracking of Speech. The model includes demographic, questionnaire, behavioural and neural predictors of reading performance in a Word Identification Fluency task using the average Mutual Information values for electrodes in the Left and in the Right Hemispheres.

**Figure 1**

*Ridge Regression Coefficient Paths Across Regularization Strengths (α): Stability of Feature Weights for Reading Prediction.*

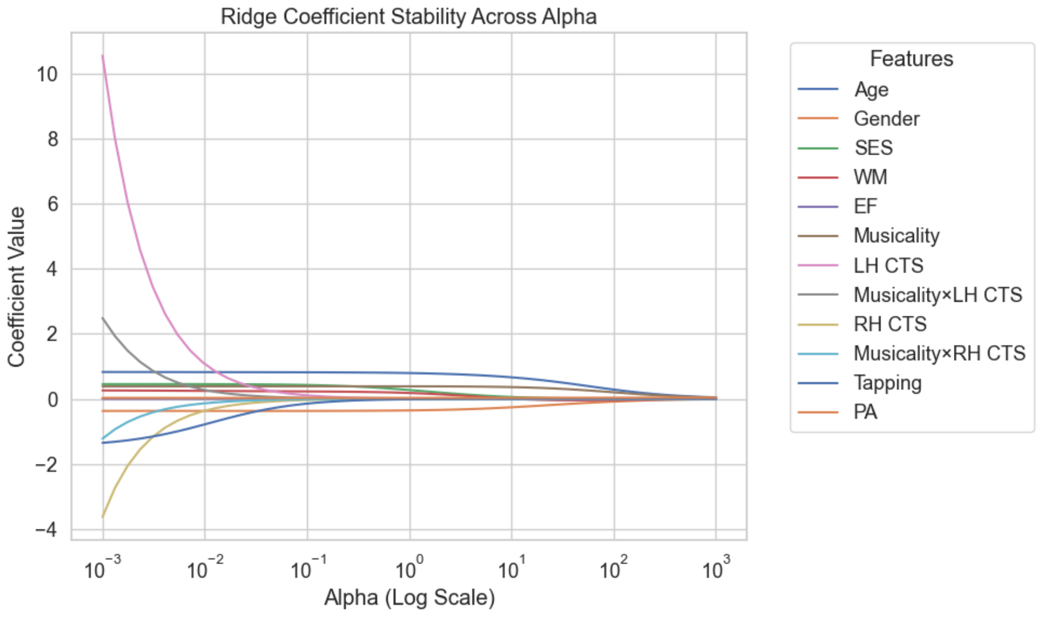


**Figure 2**

*Feature Correlation Matrix between all predictors of Reading scores.*

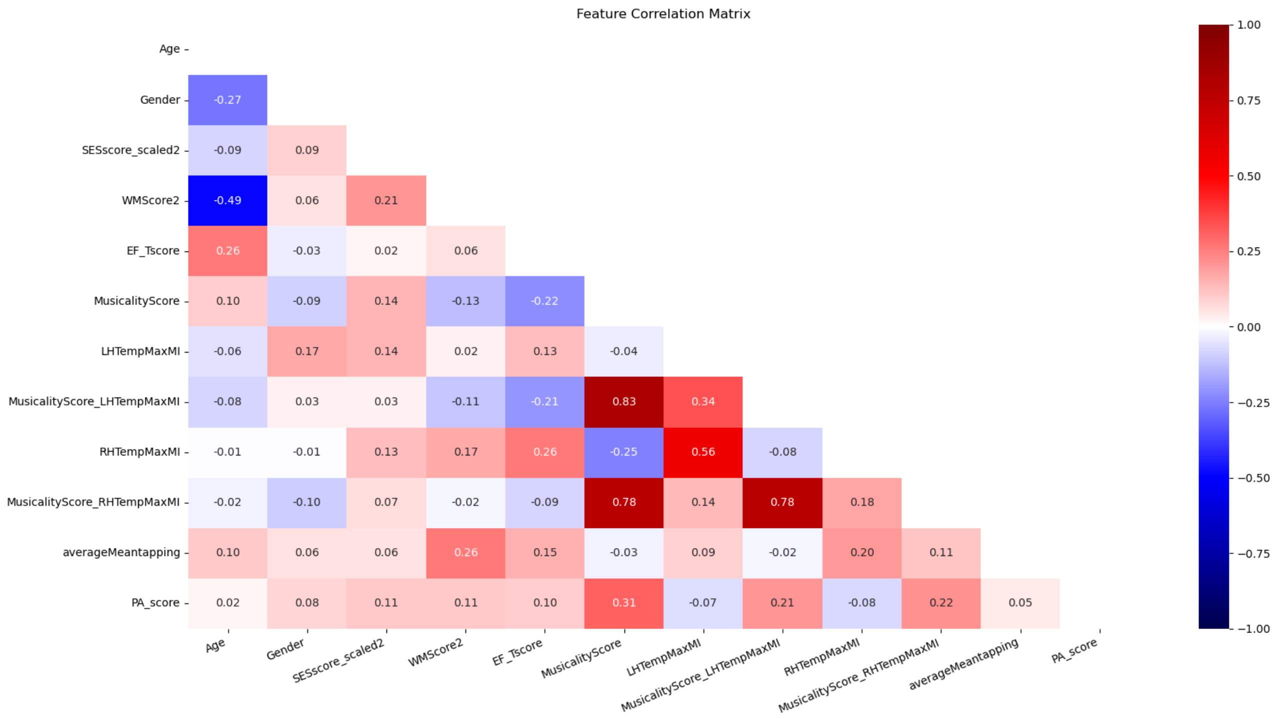

**Supplementary Materials Methods 1**

*Phonological Awareness Task - Words*

*Primary 1, 2 and 3*: rich, hawk, passed, roof, sea, shout, arms, smile, fish, woof, his, ling, life, patch, thin, when, add, learn, chips, taste, wrapped, dead, bull, meant.

*Primary 4*: words, mind, pale, short, goat, pool, hook, his, moves, stood, wet, sit, have, knows, tape, tricks, near, through, mole, who, mail, night, why, parks.

**Supplementary Materials Methods 2**

*Word Identification Fluency Task – Words*

*Primary 1*: all, am, are, at, ate, be, black, brown, but, came, did, do, eat, four, get, good, have, he, into, like, must, new, no, now, on, our, out, please, pretty, ran, ride, saw, say, she, so, soon, that, there, they, this, too, under, want, was, well, went, what, white, who, will, with, yes

*Primary 2*: after, again, an, any, as, ask, by, could, every, fly, from, give, going, had, has, her, him, his, how, just, know, let, live, may, of, old, once, open, over, put, round, some, stop, take, thank, them, then, think, walk, were, when

*Primary 3*: always, around, because, been, before, best, both, buy, call, cold, does, don’t, fast, first, five, found, gave, goes, green, its, made, many, off, or, pull, read, right, sing, sit, sleep, tell, their, these, those, upon, us, use, very, wash, which, why, wish, work, would, write, your

*Primary 4*: about, better, bring, carry, clean, cut, done, draw, drink, eight, fall, far, full, got, grow, hold, hot, hurt, if, keep, kind, laugh, light, long, much, myself, never, only, own, pick, seven, shall, show, six, small, start, ten, today, together, try, warm
